# Structural enrichment attenuates colitis-associated colon cancer

**DOI:** 10.1101/2024.02.13.580099

**Authors:** Delawrence J. Sykes, Sumeet Solanki, Sahiti Chukkapalli, Keyonna Williams, Erika A. Newman, Kenneth Resnicow, Yatrik M Shah

**Author notes:** **Corresponding Author’s email address:** and.

## Abstract

Colorectal cancer (CRC) is a major public health concern and disproportionately impacts racial/ethnic minority populations in the US. Animal models are helpful in examining human health disparities because many stress-induced human health conditions can be recapitulated using mouse models. Azoxymethane (AOM)/ dextran sodium sulfate (DSS) treatment can be used to model colitis-associated cancers. While colitis-associated cancers account for only 2% of colon cancers, the AOM/DSS model is useful for examining links between inflammation, immunity, and colon cancer. Mice were housed in enriched and impoverished environments for 1-month prior to behavioral testing. Following behavioral testing the mice were subjected to the AOM/DSS model. While our analysis revealed no significant behavioral variances between the impoverished and enriched housing conditions, we found significant effects in tumorigenesis. Enriched mice had fewer tumors and smaller tumor volumes compared to impoverished mice. African Americans are at higher risk for early onset colorectal cancers in part due to social economic status. Furthermore, housing conditions and environment may reflect social economic status. Research aimed at understanding links between social economic status and colorectal cancer progression is important for eliminating disparities in health outcomes.

## Introduction

Colorectal cancer (CRC) is the 4^th^ leading cause of cancer incidence, ranking 3^rd^ in mortality in the US [1-3]. CRC has a 91% survival rate when detected at stage I; however, by stage IV the survival rate drops to 11%. Patients with inflammatory bowel disease (IBD) are at greater risk of developing colon cancer [4]. Colitis associated cancers account for 2% of CRCs, however, there is a strong emphasis for studying colitis associated cancers due to links between IBD, inflammation and CRC [4]. Studying the relationship between stress and colitis associated cancers is an important avenue for future research concerning cancer health disparities[5-7].

Cancer health disparities represent a persistent health problem among racial/ethnic minorities and rural populations in the US [8]. Blacks and African Americans experience the highest mortality rates across most leading cancers including CRC. Determinants of excess cancer risk include poor access to treatment, underutilization of screening [9, 10], inherited risk, and more aggressive forms of the disease. Social stress may also be a factor in colon cancer health disparities [11-13]. Colorectal cancers disproportionally impact impoverished individuals and those from historically minoritized communities [14, 15]. Social and environmental stress are linked to colon cancer in humans and animals [16-18]. The causal pathways between poverty/race/discrimination and disease outcomes include poor diet, physical inactivity, obesity, lower screening rates, altered microbiome, compromised immunology, and epigenetic processes [19, 20].

Animal models can be helpful in elucidating human health disparities. Many stress-inducing conditions experienced by humans can be recapitulated using mouse models. For example, mouse protocols for overcrowding, learned-helplessness, social isolation and impoverished environments have been developed and validated [21-24]. Animal models, despite some limitations, have several advantages over human models. First, humans experience chronic stress over a period of years whereas mice can be exposed to chronic stress conditions for days or weeks. Humans may take years to exhibit deleterious effects, whereas, in mice marked impact on psychosocial behavior can be observed in weeks or months. Second, human studies also require relatively large sample sizes to detect links between stress and disease, in part due to rarity of cancer development. Third, manipulations that require noxious stimuli such as overcrowding, learned-helplessness, social isolation and impoverished environment exposures are often unethical to conduct using human subjects. Lastly, mouse models offer genetic knockouts and tight control over onset of disease and disease progression which can increase the incidence and severity of disease and thereby enhance the ability to detect health effects of stress models.

Psychosocial stress can increase risk for several cancers, including colon cancer [25-27]. However, the precise mechanisms by which stress impacts tumor initiation and progression are still unknown. Recent animal studies investigating the correlation between chronic mild stress and its impact on cancers have mixed outcomes [28-30]. While in certain instances stress induction appears to significantly affect tumorigenesis, in other scenarios, it shows no discernible effect at all [31]. Structural enrichment of living condition in animals appears to have positive outcomes [32] including; enhancement of spatial memory [33], lower anxiety, increased neurogenesis, and modest weight gain [15].

Although the impact of enrichment appears beneficial, there may be confounding effects due to variation in the amount of time animals are exposed to structural enrichment; optimally 3-weeks [16]. In our study we used a colitis-associated colon cancer model and assessed the impact of impoverished or enriched housing. Azoxymethane (AOM)/ dextran sodium sulfate (DSS) model of CRC [34] that impoverished housing increased tumor number and tumor burden.

## Methods

### Colitis-associated colon cancer model

C57BL/6J mice (N=20) were obtained from Jackson Laboratories and housed at the University of Michigan on 12:12 day/night cycle and fed *ad libitum*. All studies were carried out according to the University of Michigan and NIH guidelines (protocol approval numbers: and overseen by the Unit for Laboratory Animal Medicine (ULAM).

Our stress and enrichment manipulations began shortly after the mice arrived on campus by sorting mice into their respective housing conditions. When the mice arrived in the vivarium at The University of Michigan they were transferred to their respective (enriched and impoverished housing conditions) for the duration of the study. For the colitis-associated colon tumor model, at six weeks of age mice were injected intraperitoneally with AOM (10 mg/kg). Three days following AOM injection mice were treated with DSS in their drinking water at 2% for 7 days and switched to regular water for 14 days. This cycle was repeated 2 more times. Body weights were recorded every 5 days. Mice were euthanized 14 days after the last cycle of DSS.

### Tumor analysis

Tumors were counted, measured, imaged, and Swiss-rolled for histological analysis as previously described [35]. The normal as well as tumor colon tissue was flash-frozen in liquid N2 and stored for gene expression assays.

### Housing and Enrichment

The mice were housed in standard sized rat cages with 10 mice per cage. The enriched condition included four elements. (1) **Enviro-dry**-takes on the appearance of brown paper shredding that the mice use as nesting material. (2) **Nestlets**-cotton like material that is soft and used as additional nesting material. (3) **Chew toys**-wood blocks that are used to encourage species typical chewing behavior and (4) **plastic huts** of assorted colors were used to encourage climbing, burrowing, and nesting in of the enriched mice. By contrast our impoverished housing condition received only the Enviro-dry nesting material.

### Behavior

AnyMaze software version 6.0 (Stoelting company) in conjunction with an overhead camera to live track the movements of the mice in open field maze (40cm length x 40cm width x 40cm height) were utilized. Behaviors were measured over a 10-minute duration pre-injection. We measured the **Number of center entries-**the number of times the animal entered the center grid of the arena. **Latency to move-**the amount of time it took for the animal to move after being placed in the corner of the arena. **Distance moved-** the total distance the animal traveled in cm for the duration of the test and **Latency to exit a dark chamber-**the amount of time it took for the animal to exit the dark box after being placed inside the box with the exit door open. The dark box was created in the facility with the following specifications: A wooden (10 cm length x 5 cm width x 5 cm height) box with a hinged lid for easy opening and closing, painted black with matte paint. We anticipated that unenriched housed mice would have fewer center entries, longer latencies until first movement, and shorter distance moved with longer latencies to exit a dark chamber, all consistent with behavior observed in more anxious mice.

### Statistics

Analyses were performed using Graphpad Prism 9.5.1. We compared means between enriched and impoverished groups using an unpaired Welch’s t-test. All error bars represent standard errors of the mean unless otherwise noted. A p-value less than 0.05 was considered significant and all group numbers and explanation of significant values are presented in the figure legends.

## Results

Mice were housed in either enriched or impoverished conditions for a month preceding the behavioral tests (Figure 1). Initial analysis revealed no significant behavioral variances between the impoverished and enriched housing conditions on behavioral outcomes. Specifically, no noteworthy effects of housing were observed on activity (Figure 2a), center entries (Figure 2b), latency until first movement in the open field arena (Figure 2c), or latency to exit the dark box into the light (Figure 2d).

**Figure 1.**
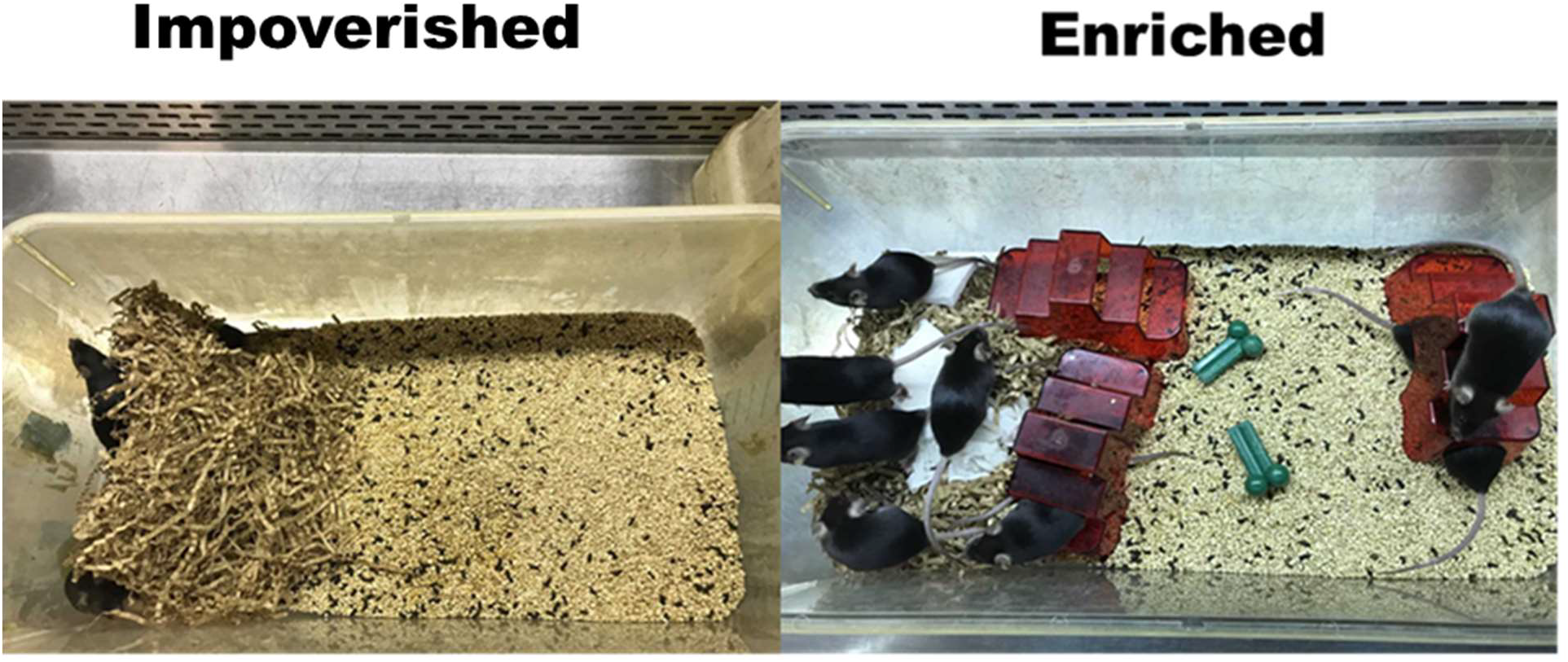
Images of (A) enriched housing versus (B) impoverished housing. Represented in image (A) are types of enrichment including *chew toys, huts, nestlets, envirodry*. Represented in image (B) is the minimum allowable enrichment for keeping mice under experimental conditions this includes, *envirodry*.

**Figure 2.**
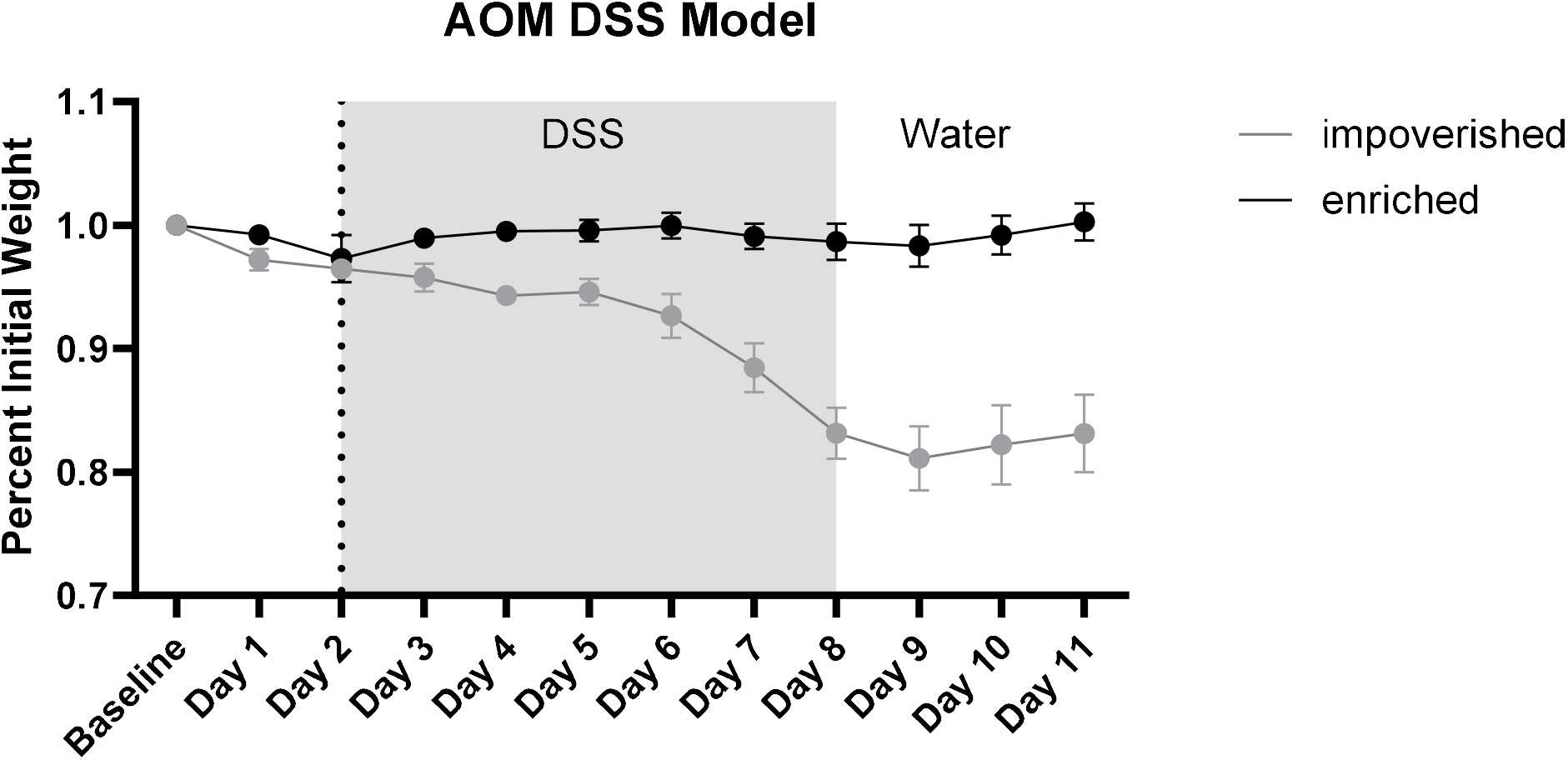
Percent initial weight and Day. Impoverished housed mice show a significant drop in weight when compared to enriched housed mice by Day8. We compare the mean weight of each group using a Welch’s t test * P<0.05.

However, impoverished housed mice lost more weight within the first 12 days following DSS administration (Figure 3). The correlation between weight loss and tumor number was r = -0.75. P<0.001

**Figure 3.**
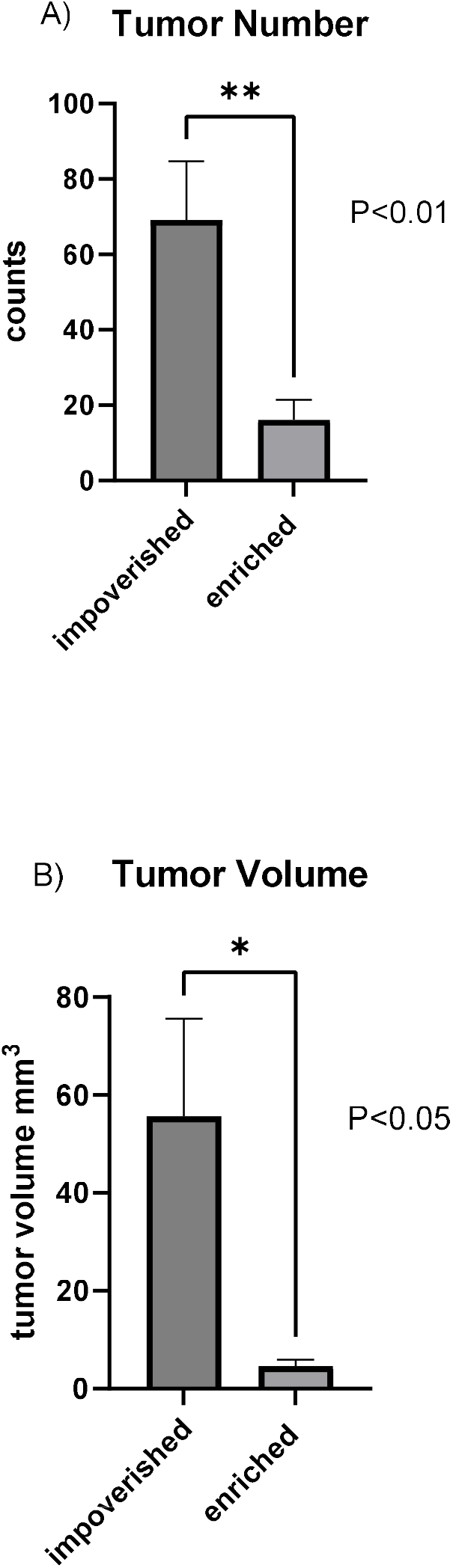
Tumor assessment (A) tumor number and (B) tumor volume. We calculated an average for each treatment and compared them using Welch’s t test. * P<0,05, ** P<0.01.

Following three cycles of DSS treatment and subsequent water recovery, the mice were euthanized, and their colons were excised. Analysis revealed that impoverished housed mice exhibited significantly (p =0.002) higher tumor number (Figure 4a) and larger tumor volumes (p= 0.011) (Figure 4b) compared to mice housed in the enriched environment.

**Figure 4.**
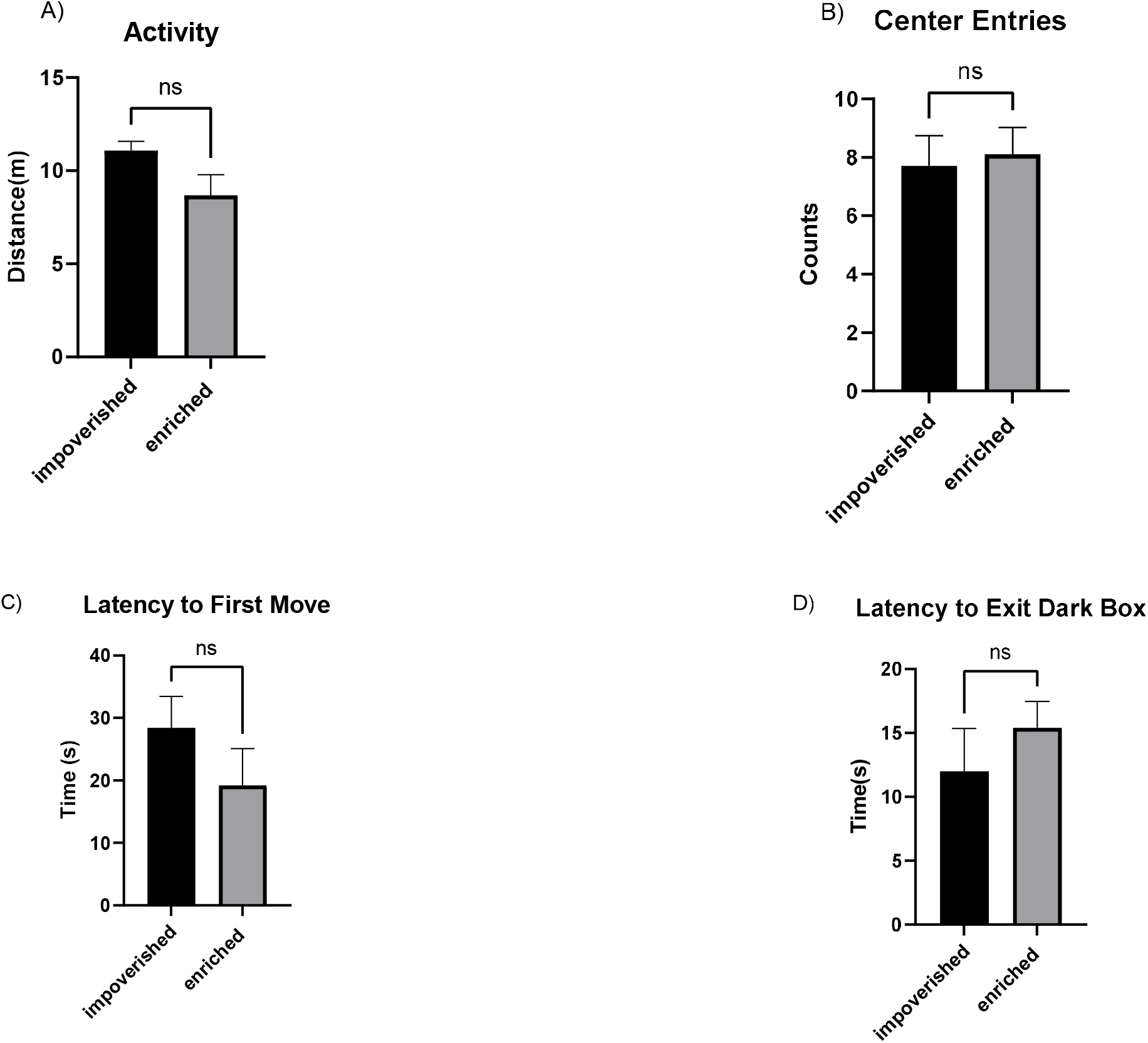
Behavioral measures calculated using the open field arena (40x 40x 40 cm). AnyMaze software was used to track the movements of each mouse in an open field. (A) Activity (B) center entries (C) latency to Move (D) latency to exit the dark box. We calculated means for each treatment group and compared them using a Welch’s t-test. NS = Not Significant.

## Discussion

We employed the open field test to evaluate anxiety-like behavior in mice following a month-long exposure to enriched or impoverished housing conditions. Surprisingly, our analysis revealed no discernible differences in anxiety-related metrics, including center entries, latency to move, latency to exit the dark chamber, or total distance moved. These behavioral outcomes have previously been observed in other stressed conditions in mice. However, it is important to note the limitations of our study, such as the brief duration of the test and the limited number of anxiety-related metrics assessed, might have limited our ability to detect behavioral effects. Notably, while some studies suggest that a 5-minute duration for the open field assay is sufficient to observe behavioral impacts, some have found longer observational periods may be needed [36].

Despite the absence of behavioral differences, we observed a striking impact of housing enrichment on tumor burden and tumor count. Mice housed in enriched environments exhibited significantly lower tumor burden and fewer tumors compared to conventionally housed counterparts. Additionally, our findings suggest that weight loss during the initial seven days of DSS administration predicts a more severe tumorigenic outcome in conventionally housed mice compared to those in enriched housing. Correlation of weight loss and tumor was r = -0.75, p =0.0007.

Currently, the mechanisms driving the observed stress-cancer relationship are not well understood. However, we hypothesize three potential explanations. First, the involvement of the central nervous system likely contributes to heightened tumorigenesis in stressed mice by communicating with the adrenals[37]. Glucocorticoids are lipid hormones that are secreted to modulate the stress response and immune regulation[38-40]. Reduced intestinal glucocorticoid production is associated with higher inflammation, thus promoting tumorigenesis[40]. Tumor cells can secrete extra-adrenal glucocorticoids inhibiting activation of T lymphocytes, thus contributing to tumor cell immune evasion[40]. Secondly, potential crosstalk between the central nervous system and the enteric nervous system might directly impact gut tumors[41]. Propranolol as a β1 and β2 adrenergic receptor blocker inhibits AOM DSS induced tumor development [42]. Similarly, cancers cells can have β adrenergic receptors and respond to the presence of catecholamines produced during stress. Thus, psychological stressors promote angiogenesis and metastasis which can be reversed by administration of β blockers like propranolol [43]. Lastly, the gut-brain axis, crucial in many cancers, is implicated in tumorigenesis. Stress triggers the release of neurohormones, including catecholamines and glucocorticoids, which have been associated with immunosuppressive effects and angiogenesis promotion, ultimately favoring cancer progression[44-46]. The link between gut dysbiosis and colorectal cancer is well established[47-50]. Microbiota from CRC patients is sufficient to induce colorectal cancer in germfree mice in the absence of carcinogen (Azoxymethane) by activating inflammatory pathways that promote intestinal tumorigenesis[51].

Notably, the bidirectional communication between the central nervous system and the enteric nervous system could hold significant implications for cancer progression in stressed mice[52]. Moreover, considering the integral role of the gut microbiome in the gut-brain axis, microbial metabolites influencing mood and neurotransmitter systems, such as serotonin, might contribute to cancer outcomes[53, 54]. Antidepressants such as Selective Serotonin Reuptake Inhibitors have shown potential in improving cancer outcomes, although current evidence regarding their effects on cancer remains inconclusive[55, 56].

In future studies, we aim to assess whether dysregulation of amino acid sensing in impoverish housed mice may link to the mTORC1 pathway [57]. Further studies are imperative to elucidate the specific mechanisms directly involved in modulating tumorigenesis in the AOM/DSS model.

There is an increase in colorectal cancer incidence among African Americans under 50 years of age compared to Caucasians under 50 [58, 59]. Thus, African Americans are at higher risk for early onset colorectal cancers, and are more likely to be diagnosed later with more aggressive forms of the disease [58, 60]. African Americans are at higher risk for early onset colorectal cancers in part due to social economic status, which can be viewed as a form of impoverished living conditions. Lower social economic status is associated with higher cancer risk including colitis associated cancers. Similarly, major depression appears more frequently among individuals with lower social economic status, especially among children and women [61]. Social economic status is a major predictor of many health and illness outcomes, yet, attention to this area of public health is in decline [62, 63]. Continued research aimed at understanding the connection between social economic status and health is an essential element of public health initiatives [64].

## Acknowledgments

This work was funded by NIH grants: R01CA148828, R01CA245546, and R01DK095201 (Y.M.S); UMCCC Core Grant P30CA046592 (Y.M.S).

## Authorship

Contribution: DS, KR and YS conceived and designed the study; DS, SS, SC developed the methodologies; DS, SS, and CS acquired the data; DS, YS, KR, EN analyzed and interpreted the data. All authors edited and provided inputs to the manuscript.

